# Effects of breast fibroepithelial tumor associated retinoic acid receptor alpha ligand binding domain mutations on receptor function and retinoid signaling

**DOI:** 10.1101/2022.12.01.518133

**Authors:** Xi Xiao Huang, Ley Moy Ng, Po-Hsien Lee, Peiyong Guan, Mun Juinn Chow, Aisyah Binte Mohamed Bashir, Meina Lau, Kenric Yi Shu Tan, Zhimei Li, Jason Yongsheng Chan, Jing Han Hong, Sheng Rong Ng, Hsiang Ling Teo, Daniela Rhodes, Patrick Tan, Puay Hoon Tan, Donald P. McDonnell, Bin Tean Teh

**Affiliations:** Cancer Science Institute of Singapore, National University of Singapore, Singapore; Cancer and Stem Cell Biology Programme, Duke-NUS Medical School, Singapore; SingHealth/Duke-NUS Institute of Precision Medicine, Singapore; Laboratory of Cancer Epigenome, Division of Medical Sciences, National Cancer Centre Singapore; Division of Medical Oncology, National Cancer Centre Singapore, Singapore; Cancer Discovery Hub, National Cancer Centre Singapore, Singapore; Oncology Academic Clinical Program, Duke-NUS Medical School, Singapore; Institute of Molecular and Cell Biology, A*STAR, Singapore; NTU Institute of Structural Biology, Nanyang Technological University, Singapore; Genome Institute of Singapore, A*STAR, Singapore; Division of Pathology, Singapore General Hospital, Singapore; Duke University School of Medicine, Durham, NC, U.S.A.

## Abstract

Point mutations in the ligand binding domain of retinoic acid receptor alpha (RARα) have been implicated in breast fibroepithelial tumors development. However, their role in the tumorigenesis of solid tumors is currently unknown. In this study, using a combination of biochemical and cellular assays, we evaluated the functional consequences of known tumor associated RARα mutations on retinoic acid signaling. All of the clinically associated mutants tested showed diminished transcriptional activities compared to wild type RARα. These mutants also exhibited a dominant negative effect, an activity which has previously been linked to developmental defects and tumor formation in mice. X-ray crystallography showed that mutants remain relatively intact structurally and the loss of transcriptional activity is due to altered co-activator recruitment. In agreement with our biochemical analyses, transcriptomics and cell growth analysis showed that the mutant RARα proteins confer resistance to growth inhibition in the presence of its ligand in phyllodes tumor cells. Although the mutations impair the receptor responses to retinoic acid, certain mutant RARα are partially reactivatable with alternative synthetic agonists. Our data provide insights into the mechanisms by which RARα mutations impact tumorigenesis.

## Introduction

Breast fibroepithelial tumors (BFT) are a heterogenous group of neoplasms characterized by a proliferation of both epithelial and stromal tissues (1). According to the World Health Organization (WHO) classification of breast tumors, fibroepithelial lesions range from the fibroadenomas and its various subtypes to the entire spectrum of benign and malignant phyllodes tumors (2). Although benign, fibroadenomas account for almost 50% of all biopsied breast tumors and occur most frequently in young women, typically between the ages of 20-30. Several studies have indicated that a particular subset of fibroadenomas might progress into the rare phyllodes tumors (3–5) which have recurrent potential, with the malignant phyllodes tumor harboring metastatic likelihood. There are no clear guidelines as to how BFTs should be managed, and surgery remains the standard of care. On one hand, conducting surgery on all patients with fibroepithelial lesions would place an enormous burden on health care systems; on the other, leaving indeterminate fibroadenomas untreated may pose a risk of progression into phyllodes tumors.

Due to a lack of understanding of the molecular events responsible for the pathology of fibroepithelial tumors, predicting their clinical behavior and ensuring optimal treatments remains challenging (6–8). To this end, we have previously characterized the genomic landscape of BFTs, which revealed distinct patterns of mutations across the tumor subtypes (9, 10). Recurrent *MED12* mutations (52%-73%) were found in both fibroadenomas and phyllodes tumors, supporting the hypothesis that phyllodes tumors may have progressed from the former. Besides *MED12*, mutations in the *RARA* gene were also frequently identified across the fibroadenoma-phyllodes spectrum, suggesting their importance in breast fibroepithelial tumorigenesis. *RARA* mutations were found to co-occur more frequently with *MED12* mutations; 20% of tumors with mutated *MED12* possessed *RARA* mutations while only 6% of tumors with wild-type Med12 were found to have *RARA* mutations. Compared to the benign fibroadenomas, phyllodes tumors harbored a higher mutational load (10), with additional mutations in the classical cancer associated genes such as *TP53, RB1* and *TERT*. Taken together, these data implicate *MED12* and *RARA* as the driver mutations in breast fibroepithelial tumorigenesis and highlight genetic features that are common among all subtypes.

*RARA* encodes retinoic acid receptor alpha (RARα), a nuclear receptor involved in the retinoic acid signaling pathway, an evolutionary conserved pathway known to be involved in many important biological processes and has been implicated in cancer (11). Retinoic acid receptors (RARα, RAR and RARψ) mediate the transcriptional responses to *all-trans* retinoic acid (RA), an active metabolite of vitamin A. As nuclear receptors, RARs regulate gene expression via the ligand-induced recruitment of transcriptional co-regulator complexes. While RARs have been known to regulate various homeostasis processes, its role in solid tumorigenesis has not been as well studied. Previously, *RARA* alterations have only been implicated in acute promyelocytic leukemia (APL), whereby the genetic translocation t(15;17)(q24;q21) resulting in fusion of *RARA* with *PML* is a molecular hallmark of this disease (12). Our genomic study of breast fibroepithelial tumors (9) was the first to discover high frequencies of point mutations in RARα associated with solid tumors. The emerging role of RARα mutations in tumorigenesis underlies novel mechanisms that warrants further investigation.

In this study, we set out to characterize the impact of *RARA* mutations on the receptor’s function and retinoic acid signaling. Our data revealed that clinically associated *RARA* mutations have impaired transcriptional activity that occur secondary to dysregulated ligand binding and coregulator recruitment. These properties result in dominant negative receptors that suppresses wild-type (WT) RARα activity, similar to what has been described for a dominant negative RARα that has been shown previously to disrupt RA signaling in mice and facilitate mammary tumor formation (13, 14). Patient-derived isogenic cell lines expressing mutant RARα remained unresponsive to agonist treatment and exhibit a downregulation of retinoic acid signaling, validating the results from our biochemical studies.

## Results

### Positions of clinically associated RARα mutations

The domain structure of RARα consists of a DNA binding domain which recognizes a specific retinoic acid response element (RARE) in the enhancer regions of RA target genes, and a C-terminal ligand binding domain (LBD) responsible for ligand binding, co-regulator recruitment and heterodimerization with retinoid X receptors (RXRs). Previously we reported that 17-32% of BFTs harbored *RARA* mutations, all of which are located in the LBD (9, 10). Besides in BFTs, missense mutations in the RARα LBD have been found in patients with APL who relapse while on RA treatment (15). Although these mutations are in the PML-RARα fusion protein, their relative positions within the RARα LBD overlap with RARα mutations found in breast fibroepithelial tumors **(Figure 1A)**. RARα mutations have also been reported in other diseases, but at low frequencies **(Supplemental Figure 1A)**. Nonetheless, most of the previously reported mutations are missense mutations and are localized in the RARα LBD, like those found in BFT and APL relapse cases **(Supplemental Figure 1, B and C)**. Due to their similarities, we suspected that these mutations may affect the RARα LBD similarly in the various conditions.

**Figure 1.**
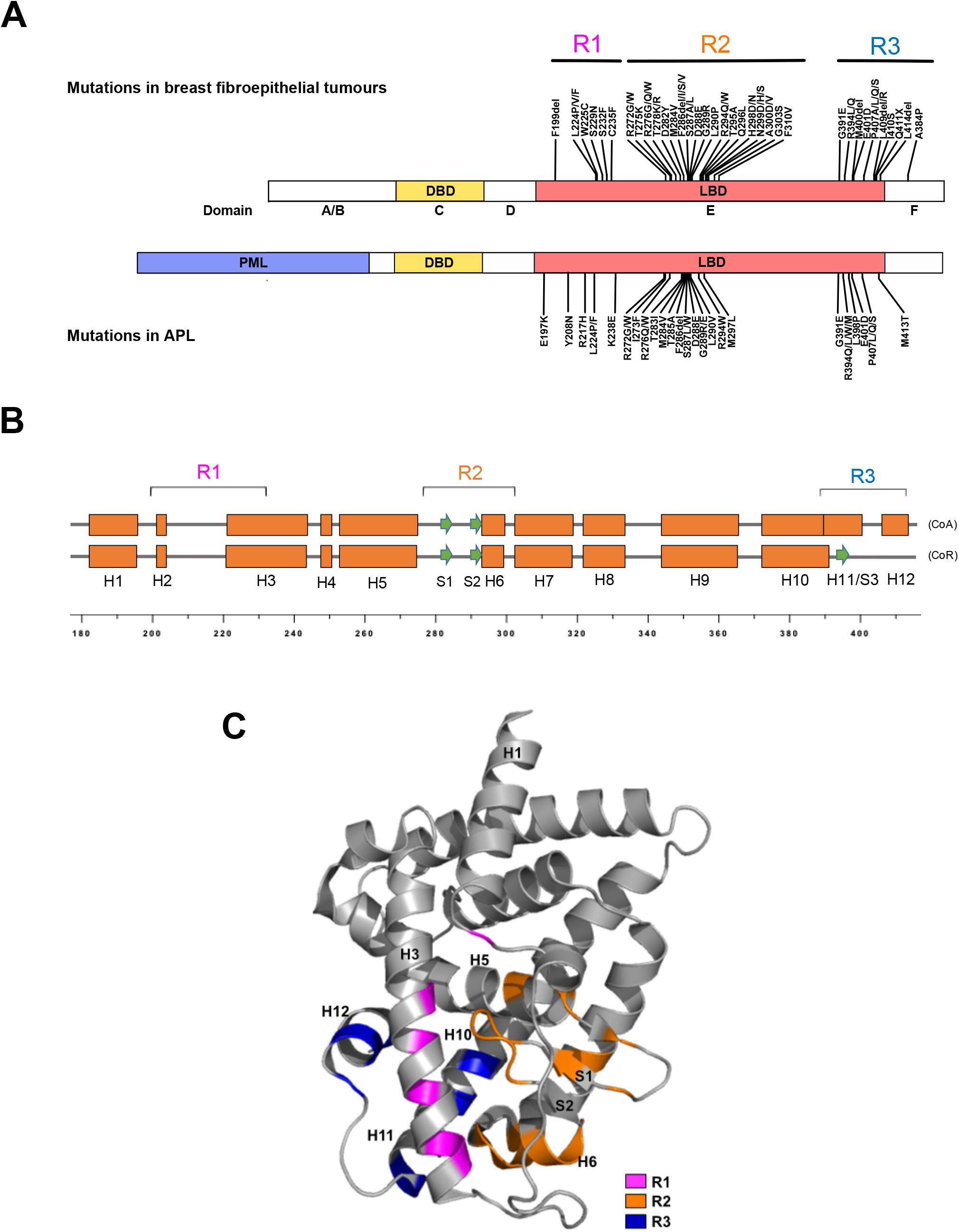
RARα mutations in diseases. (A) Domain structures of full length RARα and PML-RARα. Distribution of breast fibroepithelial tumor-associated mutations in full length RARα and RA resistant APL-associated mutations in the RARα LBD of PML-RARα. (B) The secondary structural elements of RARα LBD in the coactivator (CoA)- or corepressor (CoR)-bound states are shown as boxes and arrows representing alpha helices and beta strands respectively. Clinically associated mutations are in the indicated R1, R2 and R3 regions. (C) Locations of clinical mutations in the 3D structure, based on the published crystal structure of RARα LBD (PDB: 3KMR). Mutations in R1, R2 and R3 are colored in magenta, orange and dark blue respectively.

From the linear domain structure, the clinical mutations appear to be distributed across three regions (termed R1-R3 here) within the twelve helices of the RARα LBD **(Figure 1, A and B)**. R1 is a region from helices H1 to H3 and mutations in this region are sparsely distributed **(Figure 1C)**. Of note, many of the mutations are densely clustered in the R2 region, which spans from H5 to H6. As this region is where many ligand-interacting residues reside, we speculated that some of the mutations in this region may directly affect ligand binding. R3 is the subdomain spanning H11 and H12 (or S3) region which undergoes ligand-induced secondary structure changes and contains the coactivator/corepressor binding surface (16). Therefore, it appears that mutations in R3 could affect the recruitment of coactivators or corepressors, resulting in altered transcriptional activities. However, it was unclear how mutations in the R1 region could affect the receptor function. We therefore sought to further understand the functional effects of each of these classes of mutations and how they impacted RA signaling.

### Mutant RARα are impaired in transactivation activity and exhibit dominant negative activity

We performed biochemical studies on a larger set of RARα mutations, consisting of 21 missense mutations and two in-frame deletions that were found in BFT as well as in APL. The transcriptional activities of the mutations were assessed in two types of mammalian cell-based reporter assays. First, a RARE-dependent reporter assay (Cignal, Qiagen) that allowed us to compare the activity of the mutants to that of exogenously expressed full-length RARα **(Figure 2A)**. Second, a one-hybrid assay that we developed that evaluated the intrinsic transcriptional activity of RARα LBD-Gal4 DNA binding domain fusions **(Figure 2B)**.

**Figure 2.**
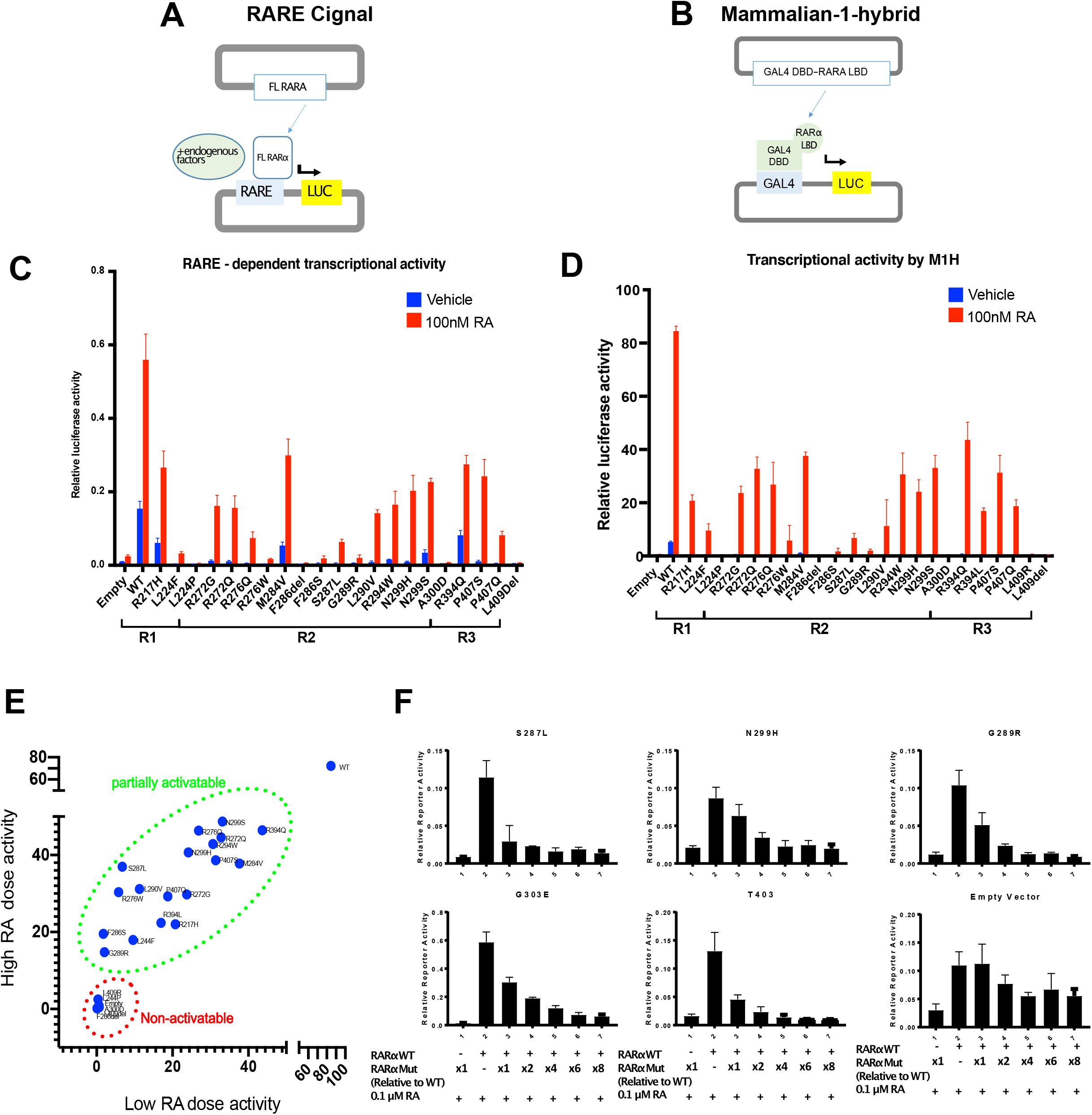
RARα mutants lose transcriptional activities and are dominant negative. (A) Schematic illustration of Cignal (Qiagen) RARE-dependent reporter assay. (B) Schematic illustration of in house one-hybrid activity assay. (C) RARE-dependent reporter assay of the transcriptional activities of overexpressed full-length WT and mutant RARα (indicated in their respective regions) in the presence and absence of 100nM RA. N=3, error bar = SD. (D) One-hybrid assay of the transcriptional activities of the wild type and mutant RARα LBD (indicated in their respective regions) in the presence and absence of 100nM RA. N=3, error bar = SD. (E) Plot of transcriptional activities of 1 μM RA treated wild type or mutant RARα vs 100nM RA treated, using combined data from Figure 2D and Supplementary Figure 2B. Tumor associated RARα mutants can be categorized into two groups: partially activatable and non-activatable. (F) RARE-dependent reporter assay of the transcriptional activities of overexpressed full-length wild type RARα, co-transfected in 293T cells with increasing amounts of mutant RARα expression plasmids, in the presence of 0.1 μM RA.

The results from the two assays were in general agreement and indicated that the basal transcriptional activities (vehicle-treated) of mutant RARα are completely abolished, except for a few mutants, such as M284V, N299S and R394Q, which exhibited significantly reduced levels of activity compared to the WT receptor **(blue bars, Figure 2, C and D)**. We titrated the input concentrations of expression plasmids to establish the maximal transcriptional activity of each mutant in the presence of 100nM RA, a physiologically relevant concentration of this hormone **(Supplementary Figure 2A)**. In this manner it was determined that all the mutants tested exhibited much lower transcriptional activities compared to WT (the maximal efficacy of the mutants relative to WT was ∼ 47% (0.7%-78%)) **(red bars, Figure 2, C and D)**. A few mutants, such as F286del, A300D and L409del, were inactive even when assayed in the presence of 1μM RA **(Supplementary Figure 2, B and C)**. These non-activatable mutants do not cluster in a specific location but are scattered across the LBD. We also tested the activities of RARα mutants with Am80, a specific synthetic agonist of RARα (RA is a pan-RAR agonist) and found the mutants to function as they did when activated with RA **(Supplementary Figure 2D)**. These results suggest that majority of the mutant RARα are weak transcriptional activators, although for some their reduced transcriptional activity can be reversed (at least partially) by increasing the dose of the agonist. Among the mutants analyzed are some that appear to be completely inactive. Thus, we have determined that tumor associated RARα mutants can be categorized into two groups: (1) non-activatable whereby no transcriptional activity can be detected even in the presence of high concentrations of RA and (2) partially activatable where the mutants exhibit transcriptional activity when challenged with at high doses of RA **(Figure 2E)**.

Of significance was the observation that some of the mutants can exert a dominant negative effect on WT RARα activity. Co-transfection of wild type RARα plasmids and RARα plasmids harboring clinically associated mutations (N299H, S287L and G289R) suppressed the RA-stimulated wild type RARα activity, as did the previously identified RARα T403 and G303E dominant negative RARα **(Figure 2F)**, confirming that the clinical mutations exhibit dominant negative activity. Nonetheless, the effect was rescued with supraphysiological concentration of the hormone (1μM) **(Supplementary Figure 2E)**.

### Disease relevant RARα mutants display dysregulated ligand binding and are defective in coregulator recruitment

Having determined that most of the disease relevant RARα mutants are expressed and that their DNA binding and RXRs binding abilities were retained, we posited that they may be defective in their ability to bind RA and/or engage coregulators **(Supplementary Figure 3, A and B)**. Thermal shift assays, using purified recombinant RARα LBDs, were performed to assess the ability of the mutants to bind RA. The thermal shift assay (TSA), or differential scanning fluorimetry (DSF), is used to perform quantitative ligand binding assays by analyzing changes in the thermal denaturation (ΔTm) properties of a protein as a function of ligand occupancy. Using this assay, we evaluated the ligand binding properties of those mutants whose function in the transcriptional assays suggested that they may have defective ligand binding properties. Surprisingly, it was determined that the that majority of mutants tested were capable of ligand binding **(Supplementary Figure 3C)**. Interestingly, the H12 mutants L409R and L409del, which were found to be transcriptionally inactive, retained ligand binding abilities, suggesting that this residue may be critical for coactivator recruitment. Indeed, using a florescence polarization assay we demonstrated that purified L409R RARα LBD proteins were incapable of interacting with a peptide from the coactivator SRC1 which reads on the integrity of the coregulator binding pocket in this receptor **(Supplementary Figure 3D)**. A mammalian two-hybrid assay was used to perform a more global assessment of the ability of the disease relevant RARα mutations to engage coregulators. Consistent with their compromised transcriptional activities, partially activatable mutants displayed reduced ability to engage the receptor interaction domains of the transcriptional coactivators, SRC2 and MED1 in the presence of physiological concentrations of RA **(Figure 3A and B)**. Interestingly the ability of mutants to interact with these co-activators increased when treated with 1μM RA; a finding that may explain why these mutants are transcriptionally active at supraphysiological levels of this hormone. Additionally, it appears that non-activatable mutants were unable to engage the interaction domains of either SRC2 or MED1, even in the presence of high retinoic acid concentration **(Figure 3C)**.

**Figure 3.**
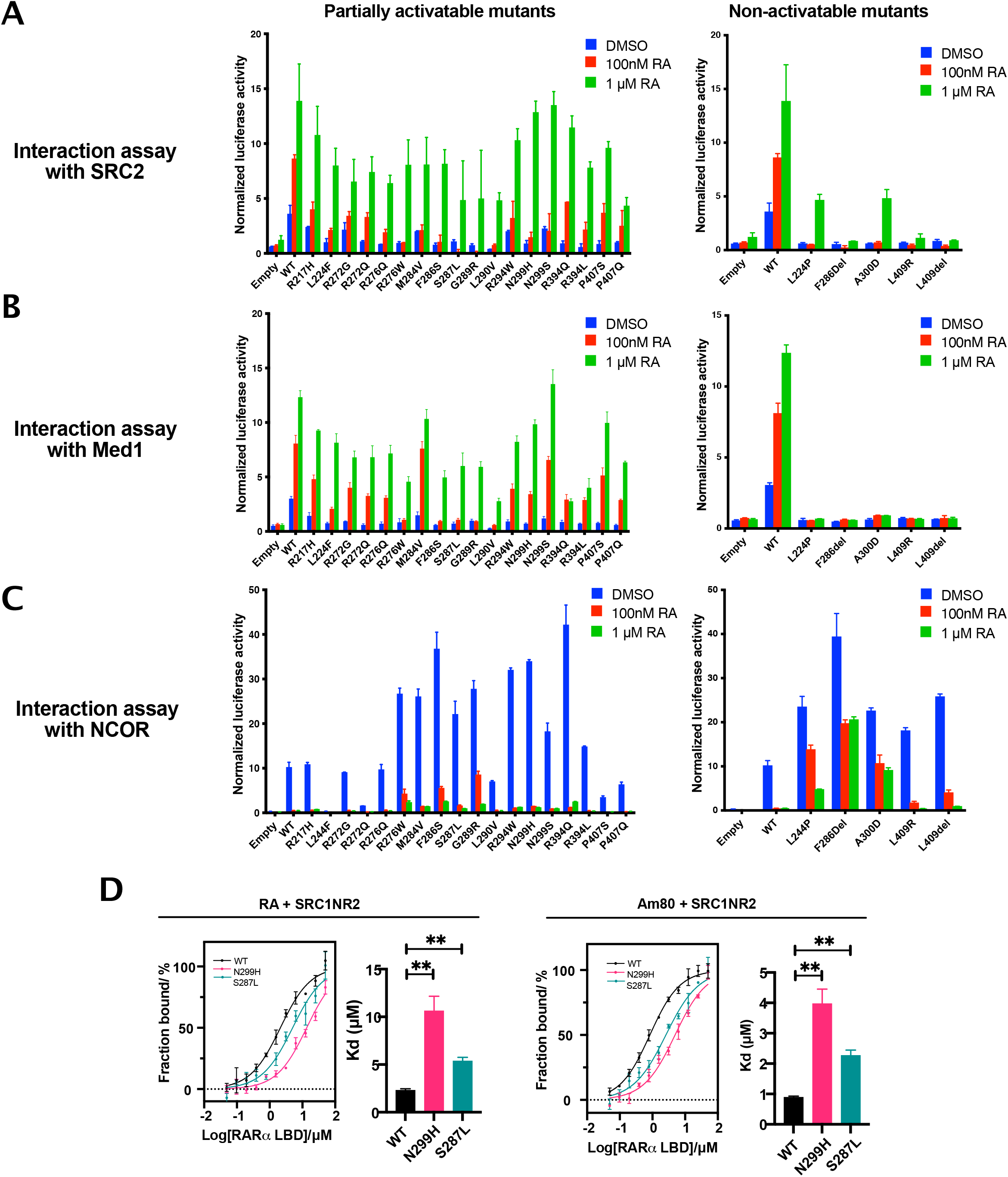
Mutant RARα possess altered binding abilities to agonists and co-regulators. Mammalian two-hybrid assay of interactions of the wild type or mutant RARα LBD with (A) fragment of coactivator SRC2, (B) MED1, a component of the transcription pre-initiation complex and (C) fragment of corepressor NCOR, treated with increasing concentrations of RA. Left panel: Partially activatable mutants, Right panel: Non-activatable mutants. N=3, Error bar = SD. (D) Binding affinities of WT or mutant RARα LBD with fragment of coactivator SRC1 (SRC1-NR2) in the presence of RA and Am80. N=3, error bar = SE. **p<0.01.

We further determined the binding affinities of two partially activatable mutants, S287L and N299H, with both co-activator and co-repressor receptor interacting peptides. Although we have observed that the mutants exhibited increased co-repressor interactions compared to WT in both basal and treated conditions in the mammalian two-hybrid assay **(Figure 3C)**, no statistically significant difference was found when we compared the binding affinities of mutant and WT RARα LBDs to co-repressor peptides (coRNR1) **(Supplementary Figure 4)**. In contrast, the binding affinities of mutant RARα and co-activator peptides were significantly lower when compared to WT in the presence of RA or Am80 **(Figure 3D)**. Taken together, these results suggest that an impaired binding ability to co-activators most likely have contributed to the low transcriptional activities of the mutants.

### Loss of H1-H3 omega loop stability affected RARα transcriptional activity

The fact that many of the clinical RARα mutations occur in residues that are unlikely to be directly involved in ligand binding or coregulator recruitment prompted us to explore further the biochemical explanation for their transcriptional defects. Our initial hypothesis was that the mutations may have altered the ligand pocket structure through rearrangements in intra-molecular interactions. We therefore used X-ray crystallography to elucidate the mutant RARα structure. We purified 20 of the clinically associated RARα mutant LBDs and screened each of the mutants under different stabilizing conditions in complexes with various ligands and either coactivator or corepressor peptides. Eventually, only one mutant (N299H) in complex with a strong agonist, Ch55, and coactivator peptide SRC1NR2, successfully formed crystals that diffracted to a resolution of 1.9Å (**Table 1**). Contrary to our initial thoughts, the overall structure of this mutant RARα is highly similar to that of the WT, with almost identical ligand pockets **(Figure 4A)**. The major structural difference identified was in the loop between H1 and H3 which is unresolved in the N299H structure but has clear electron densities in the WT structure **(Figure 4B)**. In the WT RARα, residues of this omega loop make stable contacts with N299. The loss of such contacts in the N299H mutant is due to the bulky imidazole ring of the histidine residue, which displaces and destabilizes the loop, resulting in the observed lack of electron densities **(Figure 4C)**. To understand if disruption of this loop is sufficient to affect RARα transcriptional activity, we probed an APL-associated mutation R217H. R217 lies on the omega loop and may be involved in loop stabilization **(Supplementary Figure 5A)**. Mutation of R217 to histidine recapitulated the biochemical properties of N299H, suggesting that perturbation of the omega-loop is sufficient to alter RARα activity **(Supplementary Figure 5B**).

**Figure 4.**
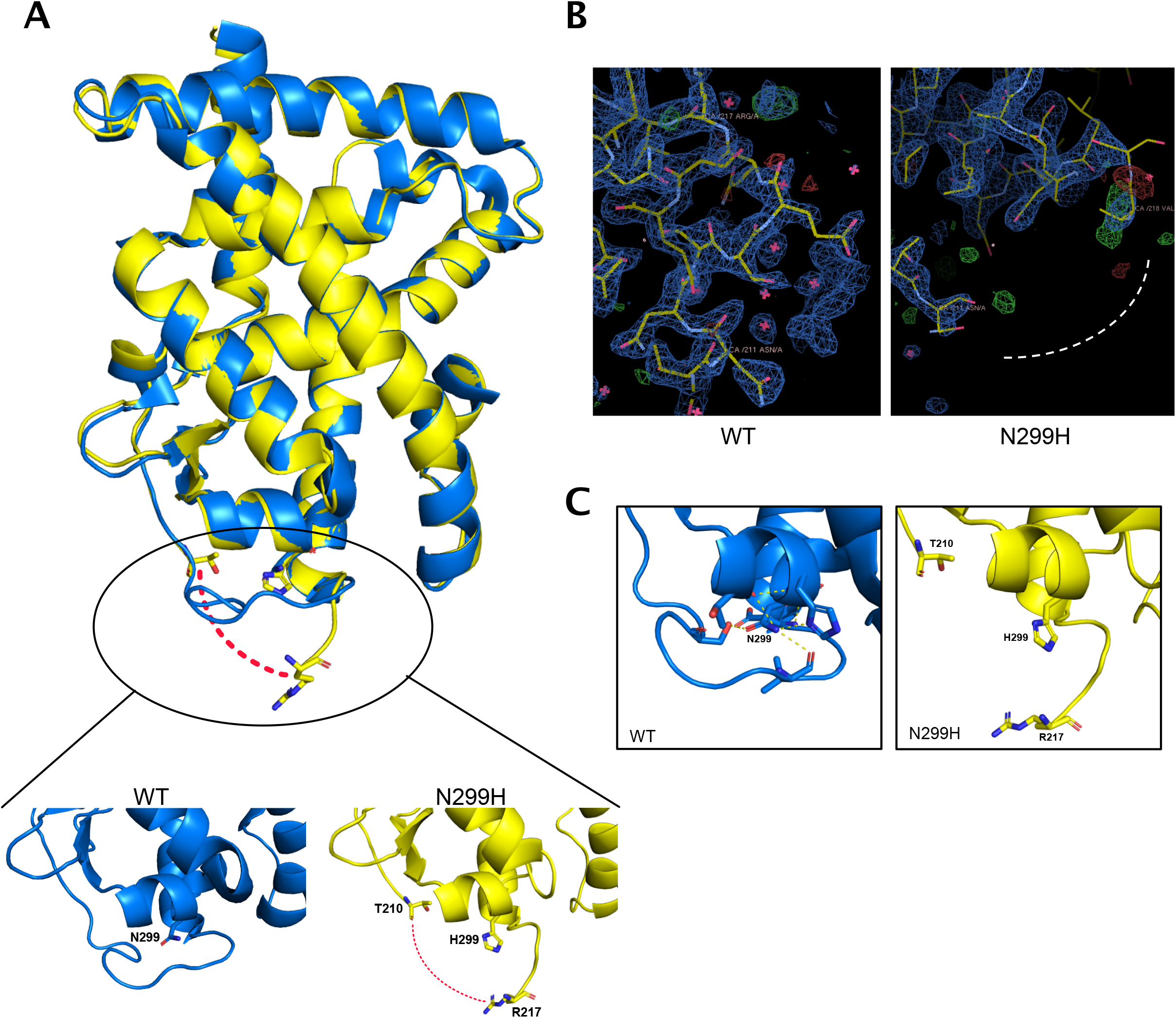
Structure of mutant RARα N299H. (A) Overlay of the overall structures of wild type RARα (PDB: 3KMR) in blue and N299H mutant in yellow. Zoomed in portions show the side chains of N299 and H299 in stick presentation. (B) Electron density maps contoured at 2, showing clear electron densities between residues 211 and 218 in the wild-type structure but missing densities in the N299H structure. (C) N299 and residues of H2-H3 loop forms hydrogen bonds (yellow dotted line), which holds the loop in position. These hydrogen bonds are lost when N299 are mutated to H299 (right).

**Table 1.**
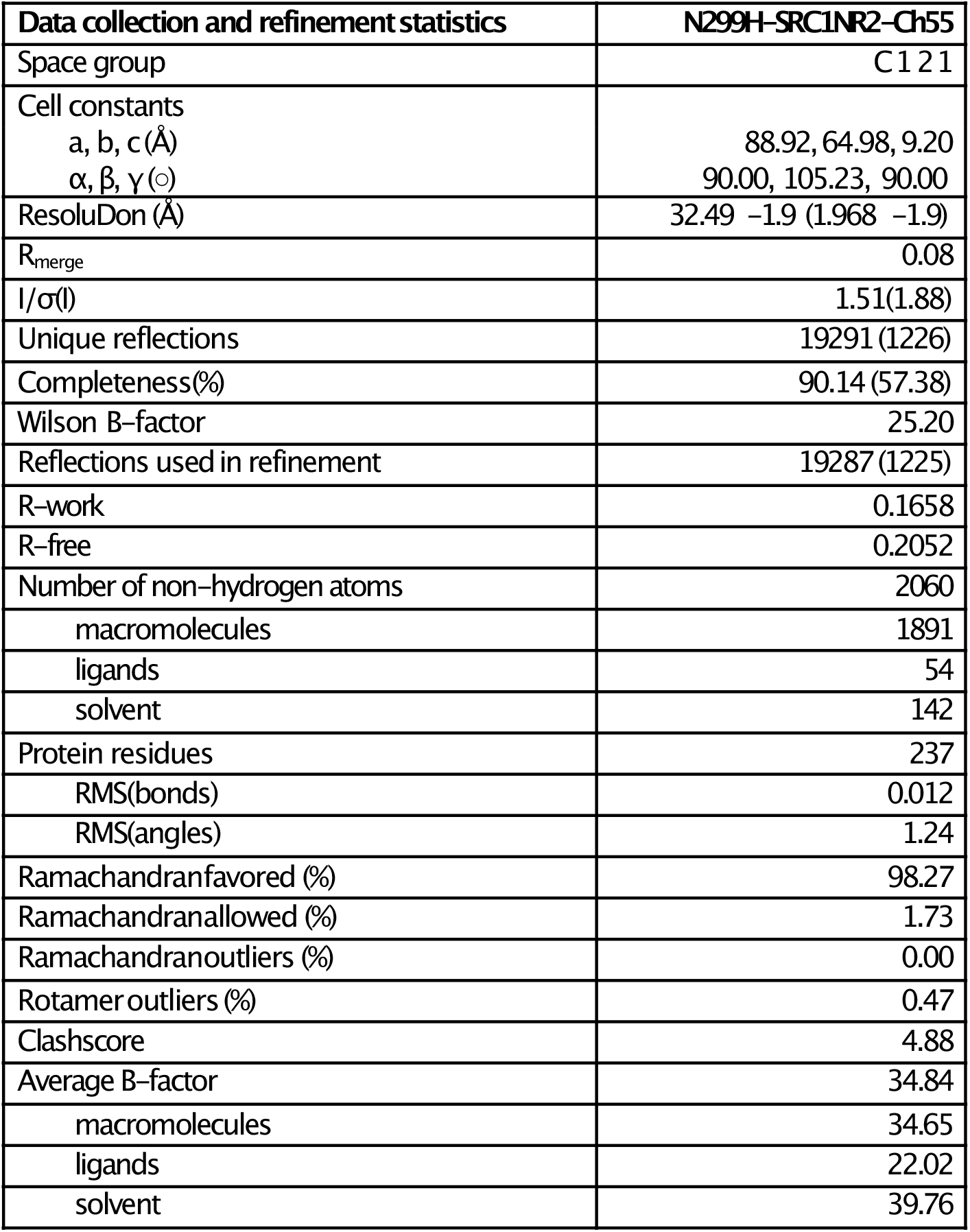
Data collection and refinement statistics of N299H-Ch55-SRC1NR2 (PDB 7WQQ)

### Sensitivity of mutant RARα to retinoids

To determine the sensitivity of cells expressing mutant RARα to retinoids, we established isogenic cell lines stably expressing doxycycline-inducible, FLAG-tagged RARα (mutant or WT) from a patient-derived phyllodes tumor cell line, PT024, which was previously established by our team **(Figure 5A)**. The parental PT024 cell line expresses mutant MED12 and WT RARα. We chose four mutations in this study: F286del, S287L (top two most common mutations found in BFT), N299H (crystal structure solved) and L409R (a H12 mutant). The growth inhibitory effects of RA and Am80 were evaluated. As expected, the growth of PT024 was inhibited by both retinoids in a dose-dependent manner (0.01-1μM). However, the cell lines expressing the RARα mutants were either less sensitive or unresponsive to retinoid treatment (**Figure 5B**). Cell cycle analysis by propidium iodide staining revealed that PT024 cells with WT RARα status (WT and control) underwent G1-S phase transition block upon Am80 treatment, as indicated by an increase in percentage cell population of G1 phase **(Figure 5C)**. In contrast, no changes were observed in the mutant RARα cell lines under the same conditions. We performed transcriptome analysis of RNAs extracted from cells expressing WT RARa and two mutants, S287L and N299H, after Am80 treatment. Gene set enrichment analysis (GSEA) revealed that retinoic acid signaling was downregulated in the mutant RARα cell lines **(Figure 5D and E)**. This was validated by the quantitative measurement of gene transcripts such as *RARB* and *CYP26A1* **(Figure 5F)**. *RARB* and *CYP26A1* are well established RAR-regulated genes commonly used as a proxy for RAR activity (17–19). Genes such as FABP5 and PDK1, which were known to be suppressed by the retinoic acid signaling pathway (20), were upregulated as expected. These results agree well with those obtained by our biochemical characterization, which concluded that mutant RARα are poor activators of retinoic acid signaling.

**Figure 5.**
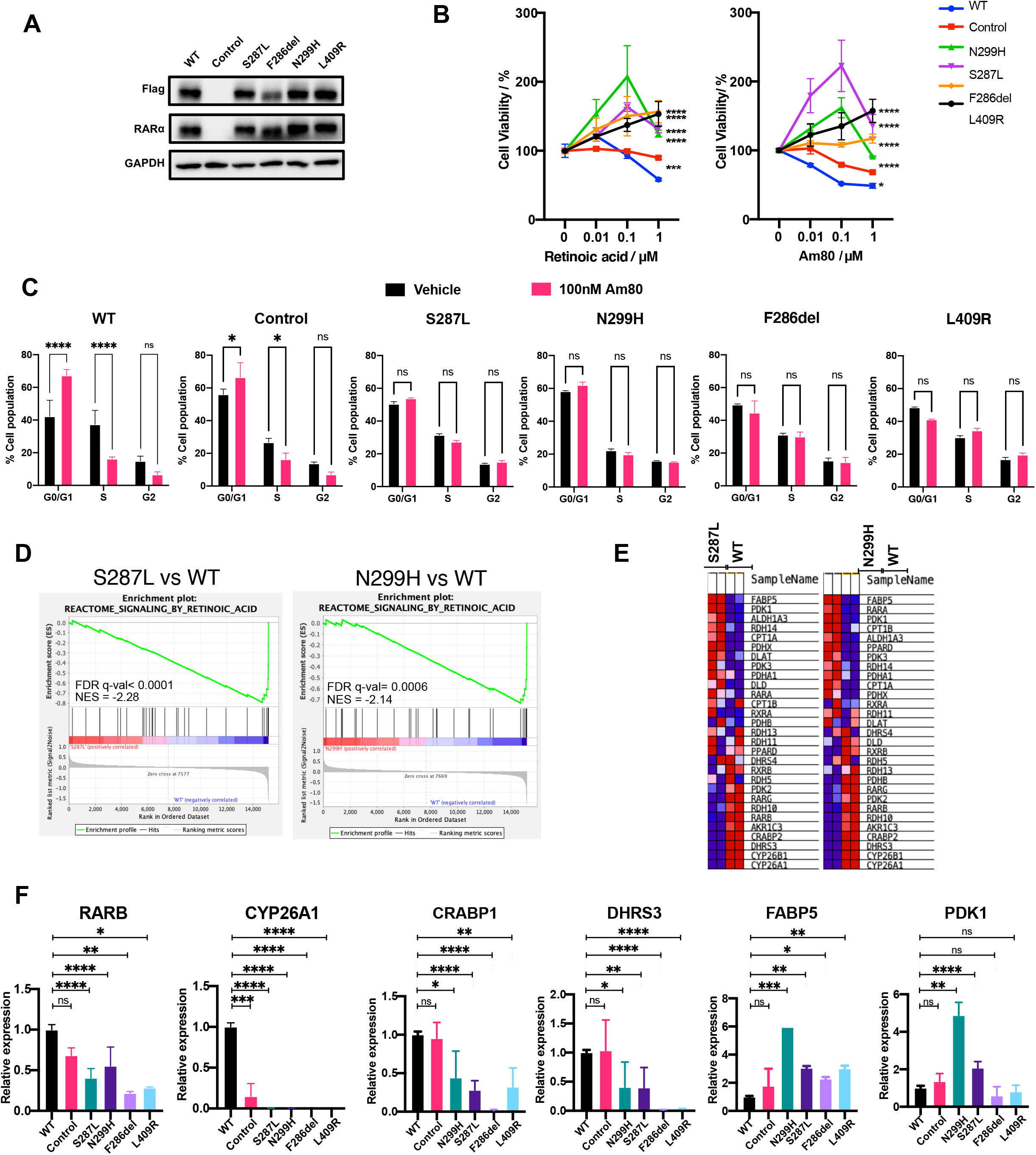
Sensitivity of mutant RARα to retinoid treatment. (A) Overexpression of Flag-tagged RARα WT and mutants in PT024 cells. (B) Cell viabilities of PT024 WT or mutant RARα expressing cells after treatment with increasing concentration of RA or Am80. N=3, error bar = SD. ***p<0.001, ****p<0.0001. (C) Cell cycle distribution of PT024 WT or mutant RARα expressing cells when treated with either vehicle or 100nM Am80. N=2, error bar = SD. *p<0.05, ****p < 0.0001. (D) GSEA analysis shows downregulation of retinoic acid signaling gene targets in mutant RARα-expressing cell lines. (E) Heatmap of genes associated with reactome retinoic acid signaling pathway (F) Quantitative PCR of retinoic acid signaling associated genes. N=3, error bar = SD. *p<0.05 **p<0.01 ***p<0.001 ****p < 0.0001.

### Targetability of mutant RARα

The fact that most of the RARα mutants show partial sensitivities to RA and retain structural integrity led us explore whether any of the mutant RARα would respond to structurally distinct, synthetic RAR agonists. Indeed, the defective transcriptional activity of mutants S287L and N299H were substantially reversed with RARα agonists **(Figure 6A)**. It was interesting however, that PT024 S287L only responded to Am580, PT024 N299H was able to respond to both Am580 and EC23 treatment, albeit slightly **(Figure 6B)**. In contrast, none of the alternative agonists tested were able to reactivate and inhibit the growth of F286del and L409R RARα expressing cells **(Figure 6, C and D)**. Although these compounds are not suitable for clinical use in patients whose tumors express these mutants (potency is too low), our data provides a proof of concept that reactivating mutant RARα signaling, particularly the partially activatable mutants, through the design and development of strong and specific ligands may be a potential therapeutic strategy in inhibiting cell growth.

**Figure 6.**
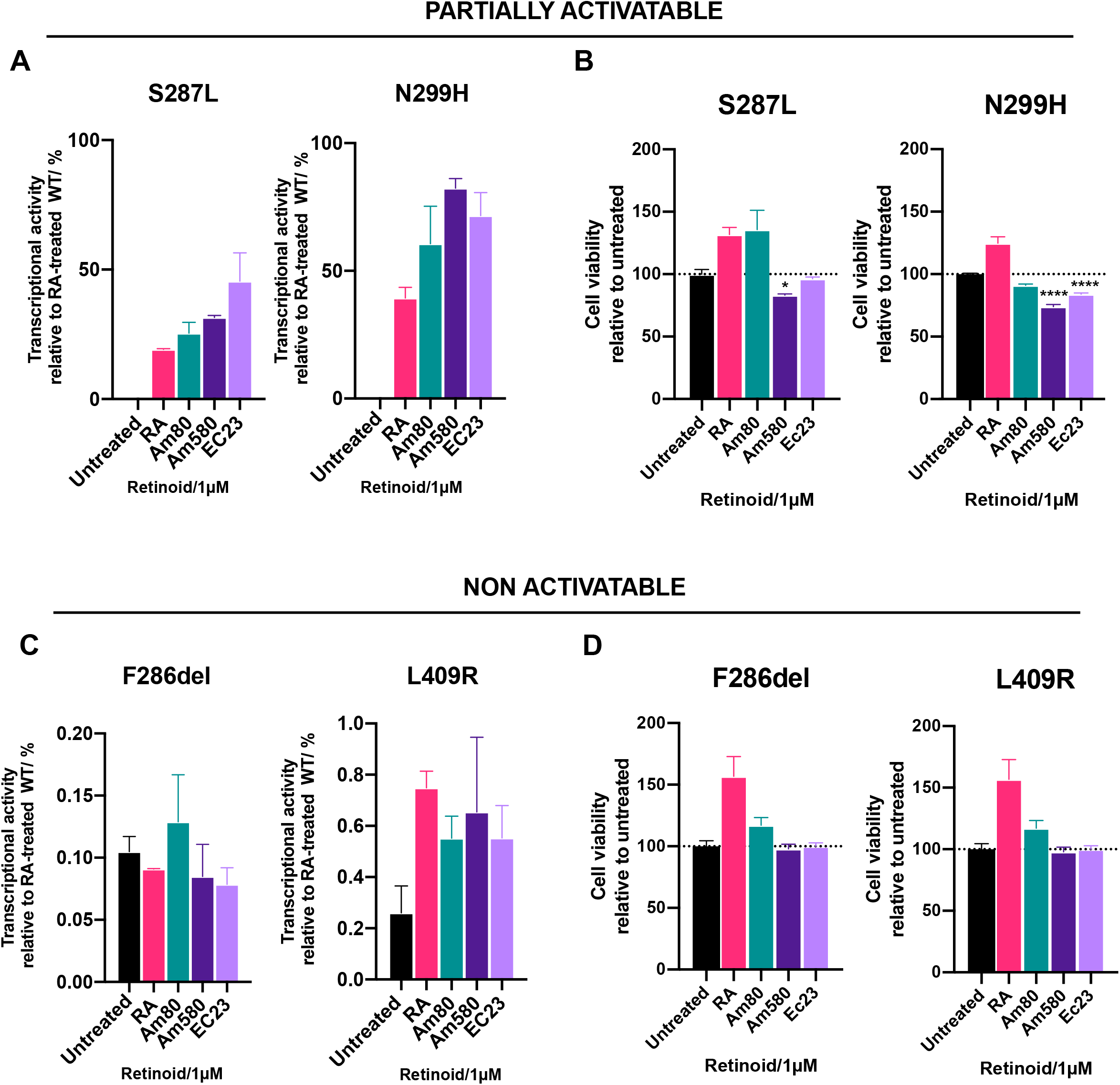
Targetability of mutant RARα. (A) Transcriptional activities (relative to RA-treated WT) of partially activatable mutants S287L and N299H after treatment with alternative RAR agonists (1μM). (B) Cell viabilities of partially activatable mutants S287L and N299H after treatment with alternative RAR agonists (1μM). N=3, error bar = SD. *p<0.5, ****p<0.0001. (C) Transcriptional activities (relative to RA-treated WT) of non-activatable mutants F286del and L409R after treatment with alternative RAR agonists (1μM). (D) Cell viabilities of non-activatable mutants F286del and L409R after treatment with alternative RAR agonists (1μM). N=3, error bar = SD.RARE RA target gene

## Discussion

Since its discovery in 1987, the role of RARα in RA signaling has been explored and described in exquisite detail and its roles, and the roles of its ligands in development, differentiation and homeostasis have been established (21, 22). In the 1980s’, the dominant negative approach was a popular technique used to study nuclear receptor function by artificially generating mutations that inactivate the signaling activities of receptors (23). Studies using dominant negative variants of RARα (C-terminal truncations or G303E mutation), have contributed significantly to our understanding of the role of RARα in disease pathogenesis (23–26). A number of studies have shown that mice expressing dominant negative variants of RARα develop leukemia, lymphoma and mammary tumors (13, 14). While the dominant negative RARα approach was initially used as a tool to dissect normal RARα physiology, our current study reveals that mutants with similar functions exist in human disease.

We have previously reported high frequencies of point mutations in RARα LBD in BFT, providing the first account of RARα mutations associated with a solid tumor. The missense RARα mutations found in BFT are remarkably like those found in RA-resistant APL relapse cases, suggesting that there are common effects induced by RARα mutations in the two different conditions. To better understand these effects, we carried out comprehensive biochemical characterizations of the most important RARα mutants. Our results show that all disease associated RARα mutants have dampened transcriptional activities, suggesting that downregulation of RARα signaling plays a role in pathogenesis. We also demonstrate that the lack of transcriptional activity is not just a simple loss of receptor integrity (ie expression), but due to altered binding affinities for ligands and to coregulators.

While it has been reported that LBD mutations in nuclear receptors tend to produce receptors that are highly repressive, such as in the case of the thyroid hormone receptors (27) and vitamin D receptor (28), we were unable to determine a change in the binding affinities of two RARα mutant receptors for co-repressors. Instead, our data show that the mutations had, in the least, affected the ability of RARα to bind co-activators; likely explaining their attenuated transcriptional activity and their dominant negative behavior. Mutations in H12, represented by the L409 mutants, were shown to completely disrupt coactivator binding. This was not unexpected as this region in RARα has been shown to be responsible for coregulator binding (29). For mutations that are found in other well less studied regions of the receptor, our structural analyses of a mutant RARα N299H revealed instability of the H1-H3 omega loop of the LBD. The importance of H1-H3 omega loop in receptor activation has been demonstrated in other nuclear receptors, specifically the thyroid receptors which belongs to the same class of nuclear receptors as the RARs and are structurally similar (30). It is believed that the H1-H3 region of the receptor crucial in the global stabilization of the LBD for ligand and co-regulator binding (31, 32). Similarly, mutations affecting this region in RARα may compromise receptor stability affecting its ability to recruit ligands or co-regulators. Using TSA, we show that the mutant LBD proteins are generally less stable (indicated by a lower melting temperature (Tm)) than the WT protein when bound to RA **(Supplementary Figure 5C)**.

Many studies have reported that RA can promote cell cycle arrest and apoptosis in various cancer cell lines (12, 33–35). A comprehensive study assessing RA-sensitivity of breast cancer cell lines has shown that the anti-tumor activity of RA is mediated by RARα (36). Hence, as expected in our established breast phyllodes tumor cell lines, we see that PT024 cells expressing WT RARα inhibited cell growth upon treatment with RAR agonists. In the case of the mutants, we have demonstrated that their expression renders cells unresponsive to RA treatment **(Figure 7)**.

**Figure 7.**
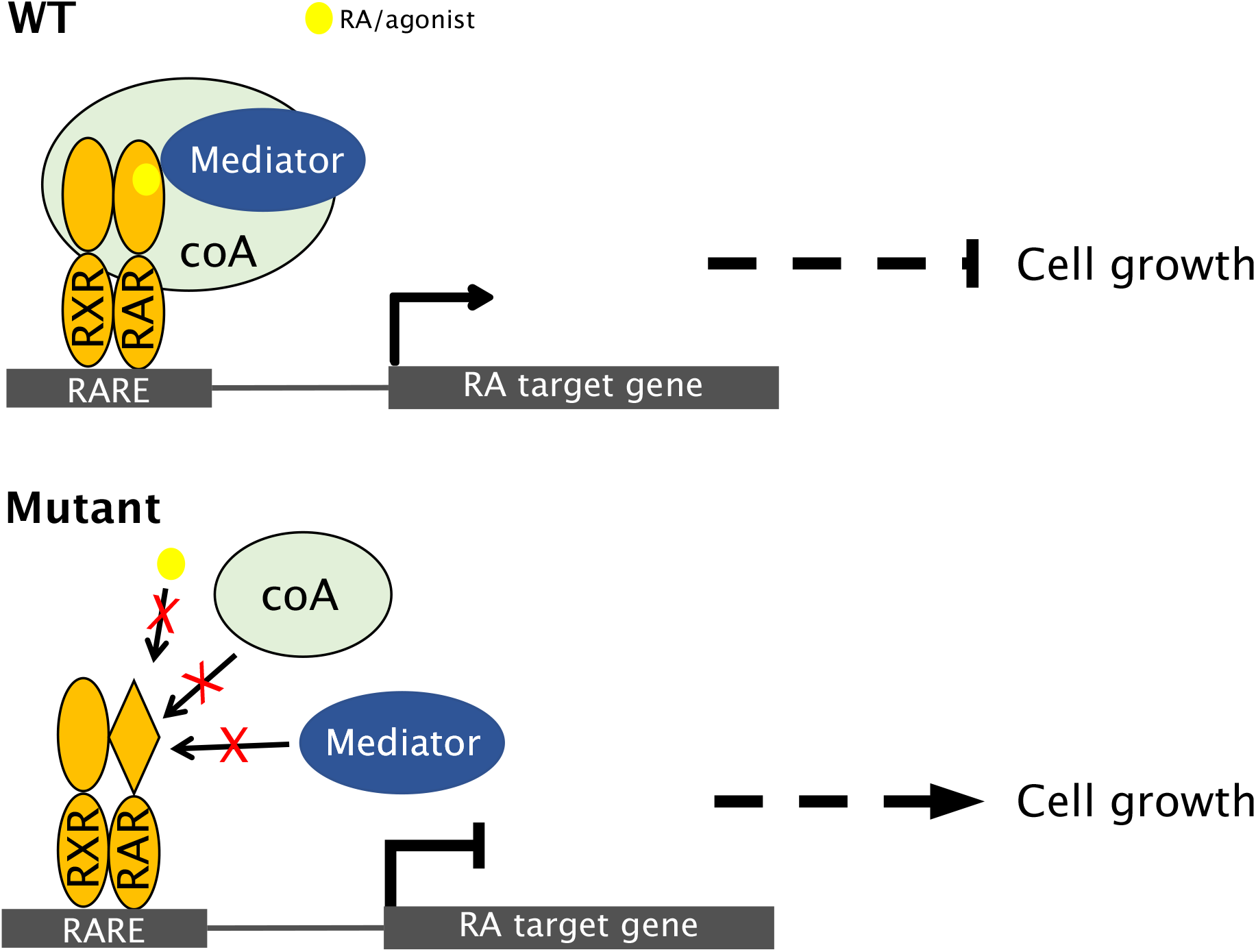
RARα LBD mutations result in altered retinoic acid signaling. Upon RA/agonist binding, WT RARα binds to co-activator (coA) such as SRC and mediator such as MED1, activating retinoic acid signaling which inhibits cell growth. Mutant RARα are poor activators of retinoic acid signaling pathway as they are defective in recruiting co-activators.

A future direction of interest will be to evaluate the potential of reactivating the signaling activities of the RARa mutants with novel ligands. Screening mutant RARα for agonists with such activities highlighted led to the identification of ligands that could activate the mutants and provide an important proof of concept that such approaches could have clinical utility. Although breast fibroepithelial tumors are mainly constituted of benign fibroadenomas, they are of huge healthcare, economic and psychological burden due to the high occurrence rate, lack of effective treatments (other than surgery) and the possibility of tumor progression. It is therefore of scientific, health and economic interest to pursue the molecular pathogenesis of breast fibroepithelial tumors and to develop effective and non-invasive therapeutics. This will also open opportunities to assess whether therapeutics targeting at mutant RARα or downstream targets of RARα will be applicable to other conditions, such as drug resistant APL relapse cases, where RARα LBD point mutations are identified.

## Methods

### Cell lines and compounds

Human embryonic kidney 293T cells were purchased from ATCC (CRL-3216). Patient derived PT024 cells were established by our team previously. Compounds tested include all-trans retinoic acid (known as RA throughout the manuscript, Sigma R2625), Am80 (aka Tamibarotene, Tocris 3507), AM580 (Tocris 0760) and Ch55 (Tocris 2020).

### RARE-dependent transcriptional activity assay

Assays were performed in 293T cells using Cignal RARE Reporter Assay Kit (Qiagen 336841) as described previously (9).

### Mammalian one-hybrid transcriptional activity assay

The sequence encoding RARα LBD (amino acids 176-421) was cloned into pBIND vector (Promega E245A) to generate pBIND-RARα LBD, which was cotransfected in 293T cells with pG5luc plasmid (Promega E249A). The transfected cells were then incubated with the indicated compounds for 24 h. Cells were then lysed and assayed for luciferase activity using dual-Luciferase Reporter Assay System (Promega E1910). Data were collected in technical triplicates and signals were normalized to Renilla luciferase readings.

### Mammalian two-hybrid assay

Assays were performed using CheckMate Mammalian Two-Hybrid System (Promega E2440). The sequence encoding MED1 (amino acids 495-699), SRC2 (amino acids 624-869), the CoRNR1 peptide region (THRLITLADHICQIITQDFARNQV) of the NCOR1 protein or the CoRNR box region containing ID1 and ID2 domains of the SMRT (amino acids 2114-2357) were cloned into the pBIND vector to generate pBIND-NCOR1 and pBIND-SMRT bait plasmids respectively. The sequence encoding RARα LBD was cloned into pACT vector to generate pACT-RARα LBD prey plasmid. Bait and prey plasmid pairs were cotransfected into 293T cells with pG5luc reporter plasmid. The transfected cells were then incubated with the indicated compounds for 24 h. Cells were then lysed and assayed for luciferase activity using dual-Luciferase Reporter Assay System (Promega E1910). Data were collected in technical triplicates and signals were normalized to Renilla luciferase readings.

### Site-Directed Mutagenesis

Mutations were introduced into RARα-encoding plasmids using Q5 Site-Directed Mutagenesis Kit (NEB E0554). Mutations were confirmed by sequencing of the resulting plasmids.

### Protein purification

The cDNA sequence encoding RARα LBD or RXRα LBD were cloned into pET His6 TEV LIC cloning vector (Addgene 29666), pET His6 MBP TEV LIC cloning vector (Addgene 29708) or pET GFP LIC cloning vector (Addgene 29716). All Addgene cloning plasmids were kind gifts from Scott Gradia. Cloned plasmids were confirmed by sequencing and transformed into BL21(DE3). Overnight transformed cultures were reinoculated into ZYP5052 autoinduction media and incubated at 37°C with shaking for 4 hours, followed by 17°C overnight. Cells were harvested and lysed by sonication. Recombinant expressed proteins were purified from lysate supernatant using standard nickel affinity chromatography followed by size exclusion chromatography.

### Thermal shift assays

TSA were performed using Protein Thermal Shift Dye Kit (Applied Biosystems 4461146) with purified WT and mutant RARα LBD proteins and the indicated compounds in five molar excess. Data were collected in technical triplicates using Applied Biosystems 7500 Fast Real-Time PCR System. Tm were analyzed using TSA-CRAFT, a software that we have developed (37).

### Fluorescence polarization

Purified H6 RARα LBD proteins was used together with FAM-conjugated SRC1NR2 or FAM-conjugated coRNR1 peptides to elucidate binding-saturation curves. FAM-SRC1NR2 was kept constant at 180nM and mixed with increasing concentrations of RARα LBD to a maximum of 50µM. RA or Am80 were added to the mixtures at a two-fold molar excess of 100µM. Reactions were incubated for 45mins at room temperature (RT) before data collection. The assay was performed in technical triplicates and mean binding affinity (Kd) was determined from three repeated experiments.

### Crystallization and structure determination

Purified N299H RARα LBD proteins were mixed with a twofold molar excess Ch55 and three-fold molar excess SRC1NR2 peptide, incubated for 1hr at 4°C and concentrated to 7mg/mL. The complex is then mixed in a ratio of 1:1 with reservoir solution containing 0.05M BisTris Propane pH 5.0, 0.05M Citric acid and 18% PEG3350. A single crystal was mounted from the mother liquor onto a cryoloop, soaked in the reservoir solution containing an additional 60% (w/v) sucrose and flash-frozen in liquid nitrogen.

Diffraction data was collected at the National Synchrotron Radiation Research Center, Taiwan, at beamline TPS 05A. Data was processed and scaled using HKL2000(38). Structure solution was performed using Phenix GUI package(39). With the package tools, molecular replacement was carried out using Phaser, with the wild type structure (PBD 3KMR, ligand removed) as a starting model. The model was built using Autobuild, followed by fitting of Ch55 into the model using LigandFit. Structural refinements were done using Coot (40) and phenix.refine (41). Atomic coordinates and structure factors for the reported crystal structure have been deposited with the Protein Data bank under accession number 7WQQ.

### Inducible cell lines generation

Doxycycline inducible RARα expressing phyllodes tumor patient-derived cells (PT024 RARα) were established with Tet-On 3G system from Takara. Coding sequence of full-length WT RARα were cloned into a PLVX-Tre3G-IRES-GFP response vector. Mutant RARα plasmids were then generated using mutagenesis. Firstly, PT024 cells were lentivirally transduced with PLVX-TET-ON-3G regulator plasmids, and the resultant cells were selected with 650µg/mL G418. The PT024/ Tet-on clones were then transduced with PLVX-Tre3G-RARA response plasmids and selected with 1 µg/mL puromycin. Subsequently, established stable cell lines were maintained in media with 325 µg/mL G418 and 0.5µg/mL puromycin. The cells were induced with 10ng/µL doxycycline for 24h before phenotypic experiments were performed.

### ATP viability assay

Induced PT024 RARα cells were seeded in 96-well white bottom plates at 2.5 × 10^4^ cells/cm^2^. After 24hr incubation, the cells were treated with the indicated concentrations of retinoids for six days, with drug and doxycycline retreatment on day three. After treatment, ATP content was assessed using CellTiter-Glo® Luminescent Cell Viability Assay (Promega, G7571) following manufacturer’s protocol. The assays were done in technical triplicates and repeated at least three times.

### Cell cycle analysis

Cell cycle analysis was done by staining DNA with propidium iodide. Induced PT024 cells were seeded at 2.5 × 10^4^ cells /cm^2^ in six well plates and treated with either vehicle or 100nM AM80 for six days, with retreatment on day three. After treatment, cells were harvested and fixed in cold 70% ethanol overnight. Then, they were washed in PBS twice and resuspended in PI staining solution (40µg/mL propidium iodide and 20µg/mL RNAse) overnight. Cell cycle distribution data was collected by BD LSRII analyzer and analyzed using FlowJo’s cell cycle analysis platform. The assays were done in technical duplicates and repeated twice.

### Preparation of whole cell lysates and immunoblotting

Whole cell lysates were prepared in Pierce RIPA Lysis and Extraction Buffer (Thermo scientific #8990), separated by SDS-PAGE and transferred onto PVDF membrane. Protein targets were detected with primary antibodies against GAPDH (Invitrogen, #MA5-15738), Flag (Sigma-Aldrich, F3165) and RARα (Cell signaling, #62294).

### RNA isolation

PT024 RARα cells were treated as described and total RNA were extracted using Monarch Total RNA miniprep kit (NEB #T2010S) according to manufacturer’s protocol.

### Quantitative PCR

Total RNA was reverse transcribed into cDNA using iScript™ Reverse Transcription Supermix for RT-qPCR (Biorad. #1708841) following manufacturer’s protocol. Quantitative PCR was performed with Luna® Universal qPCR Master Mix (NEB #M3003) and Biorad CFX96 Real time PCR system. qPCR was performed in technical triplicates and repeated at least thrice. Results fold change values are normalized to *GAPDH*.

Primers sequences used are GAPDH: 5’ ggagcgagatccctccaaaat 3’ and 5’ ggctgtcatacttctcatgg 3’, RARB: 5’ tccgaaaagctcaccaggaaa 3’ and 5’ ggccagttcactgaatttgtcc 3’, CYP26A1: 5’ tccagaaagtgcgagaagag 3’ and 5’ tcttcagagcaacccgaaac 3’, CRABP1: 5’ atcggaaaacttcgaggaattgc 3’ and 5’ aggctcttacagggcctcc 3’, DHRS3: 5’ actgagtgccattacttcatctg 3’ and 5’ catcactgtccattaggctcttc 3’, FABP5: 5’ tgaaggagctaggagtgggaa 3’ and 5’ tgcaccatctgtaaagttgcag 3’, PDK1: 5’ ctgtgatacggatcagaaaccg 3’ and 5’ tccaccaaacaataaagagtgct 3’.

### Transcriptomic analysis

Total RNA samples were isolated as described above in duplicates and whole transcriptome sequencing were performed on Novoseq 6000 sequencing system (Novogene, Singapore). Qualities of raw FASTQ data files were assessed with MultiQC (42). Data was mapped using STAR (v2.7.3a) (43), with mapping results checked by RSeQC (44). Gene expressions were quantified using RSEM (v1.3.3) (45) and varianceStabilizingTransformation in DESeq2 was used to normalize the raw counts. Differentially expressed genes were then identified with DESeq2 (v1.24.0) (46). No batch correction was required as the RNAseq libraries were sequenced in one batch. Gene set enrichment analysis were performed using GSEA (v4.0.3) (47) on Reactome gene sets (48) and Hallmark gene sets (49). Significance of the enrichments was assessed using FDR q-value. RNAseq data have been deposited under BioProject (accession number PRJNA868864).

### Native gel assay

Reactions were set up by mixing 10 µM of GFP-RARα proteins with 50 µM or 10 µM of biotinylated-RXRα for fluorescence or biotin detection respectively. Reactions were loaded directly onto non-denaturing PAGE performed at 100 V in 4 °C. Gels were visualized by fluorescence scanning (Typhoon Trio, GE Amersham) for GFP detection followed by Coomassie blue staining for total proteins. For biotin detection, proteins were transferred onto a PVDF membrane, probed with Streptavidin Alkaline-Phosphatase (Promega V5991) and detected by colorimetric assay (Promega S3841).

### Electrophoretic mobility shift assay

The open reading frame of RARα was cloned into pCMV 3X Flag expression vector and the mutant RARα plasmids generated by mutagenesis. These plasmids were transfected into HEK293T cells and incubated for 48hrs. The cells were then harvested, and the nuclei released using a hypotonic buffer (10mM HEPES; 60mM KCl; 0.075% (v/v) NP40; 1mM EDTA; 1mM DTT; 1mM PMSF; final pH7.6). The nuclear fraction was then isolated by resuspending nuclei with a high-salt buffer (20mM Tris pH 8.0, 1.5mM MgCl2, 400mM NaCl, 0.2mM EDTA, 1mM PMSF and 25% (v/v) glycerol). Biotinylated probes containing the DR5 response elements with the sense strand sequence 5’ CAT-AAG-GGT-TCA-CCG-AAA-GTTCAC-TCG were purchased from IDT. Three micrograms of nuclear extracts were incubated in binding buffer (25mM Tris pH7.5, 80mM NaCl, 5mM MgCl_2_, 5% glycerol, 50ng/µL poly (dI.dC) and 2% NP40) with 150nM of duplex probe for 30mins at RT. For super-shift EMSA, nuclear extracts and probe were incubated for 15mins at RT after which, 1µL of Flag antibody was added and the reaction mix was incubated for an additional 15mins. The reaction mix was then resolved by native gel electrophoresis and transferred onto a nylon membrane. Biotinylated probes were detected using Chemiluminescent nucleic acid detection module (Thermo Scientific # 89880) following manufacturer’s protocol.

## Author contributions

LMN and BTT supervised the work. XXH and LMN designed experiments and interpreted data. PHL, LMN and XXH performed structural analyses. PYG performed bioinformatics analysis. XXH, LMN, ML, KYST, MJC, ABMB and ZML performed experiments. JHH, SRN, JCYS, HLT, DR, PT, PHT, DPM contributed to the study directions and provided advice. XXH and LMN wrote the manuscript with the help of JHH, DPM and BTT.

## Acknowledgments

This work was supported by National Medical Research Council [NMRC OFIRG16may055], the National Research Foundation Singapore and the Singapore Ministry of Education under its Research Centres of Excellence initiative as well as Singapore Translational Research (STaR) Investigator Award [project ID MOH-00248].

