## Supplementary figures for "Effects of breast fibroepithelial tumor associated retinoic acid receptor alpha ligand binding domain mutations on receptor function and retinoid signaling"

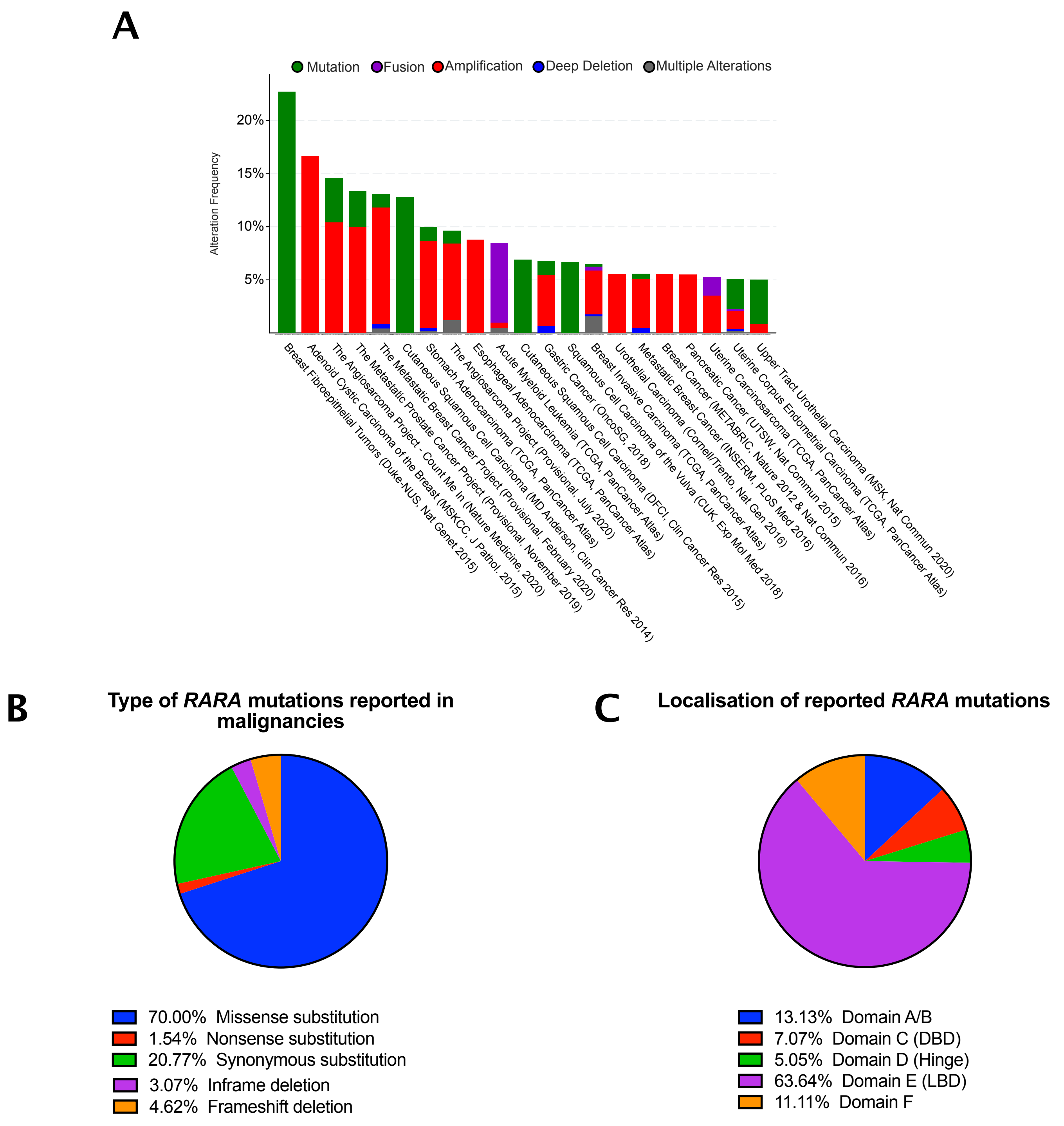


Supplementary Figure 1. RARα mutations reported in other diseases. (A) Frequencies of RARα mutations reported in various cancer studies. Data retrieved from curated set of non-redundant studies from cbioportal. (B) Pie chart of the types of RARα mutations reported in malignancies. Data retrieved from COSMIC resource database. (C) Locations of missense mutations in the RARα domain structure. Data retrieved from COSMIC resource database.

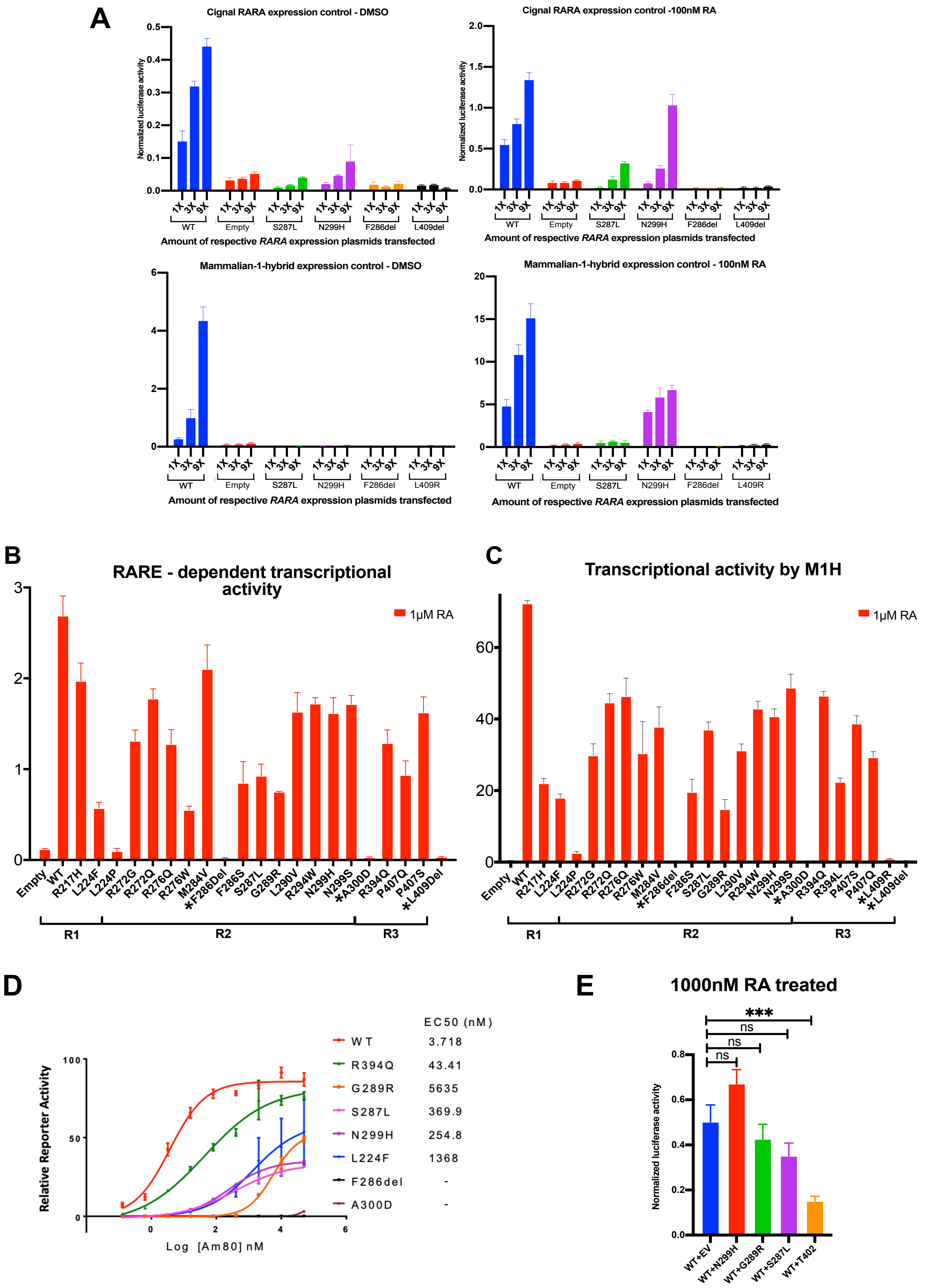


Supplementary Figure 2. Transcriptional activities of WT and mutant RARα at high RA dosage. (A) Cignal and mammalian-1-hybrid transcriptional activities of WT and mutant RARα, transfected in 293T cells in increasing amounts. (B) RARE-dependent reporter assay of the transcriptional activities of overexpressed full-length wild type and mutant RARα in the presence of 1μM RA. (C) One-hybrid assay of the transcriptional activities of the wild type and mutant in the presence of 1μM RA. (D) Efficacies of transcriptional activation of the wild type and mutant RARα LBD by the RARα-specific agonist, Am80. (E) Transcriptional activity in the presence of 1M RA after co-expression of equal amounts of WT and mutant RARα.

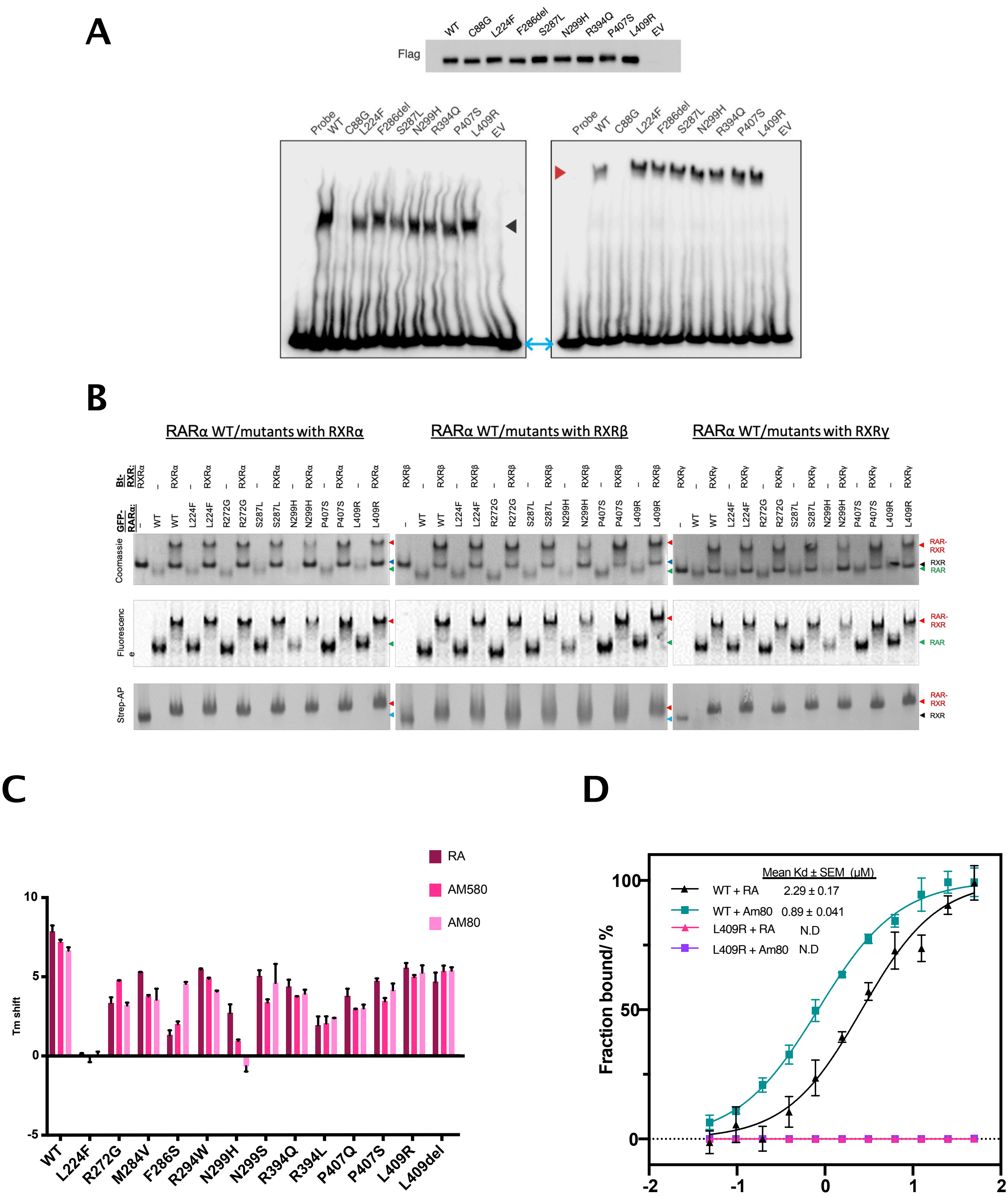


Supplementary Figure 3. DNA and RXR binding assays. (A) Flag - RARα amount were normalized in nuclear extracts and used in EMSA. EMSA of nuclear extracts with DR5 probe (left) and supershift EMSA with Flag antibodies (right). C88G, a DNA - binding mutant, was included as a control. Symbols: Black arrow – DNA shift, Red arrow – Supershift, Blue double arrow – Free probe. (B) Native gel assay of Biotin-RXRα and GFP-RARα complexes resolved by non-denaturing PAGE and visualized by Coomassie blue staining (showing total proteins), fluorescence imaging (showing GFP-tagged proteins and complexes) and streptavidin-AP blotting (showing biotin-tagged proteins and complexes). (C) Interactions of ligands with wild type or mutant RARα by TSA. N=3, error bar = SD. (D) Binding affinities of WT and mutant RARα L409R LBD with SRC1 nuclear receptor interacting fragment (SRC1-NR2) in the presence of RA and Am80.

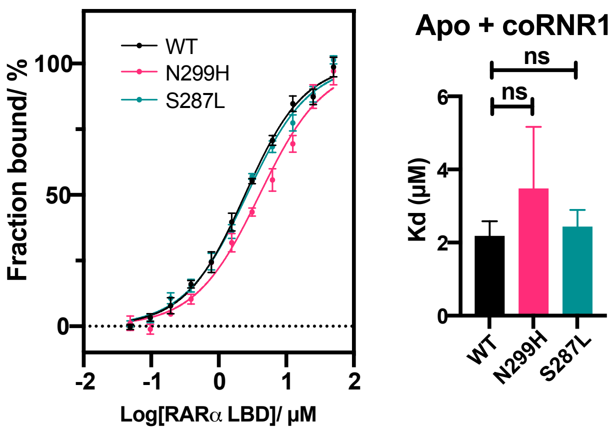


Supplementary Figure 4. Binding affinities of WT and mutant RARα N299H and S287L LBD with fragment of co-repressor NCOR1 (coRNR1) in basal conditions. N=3, error bar = SD.

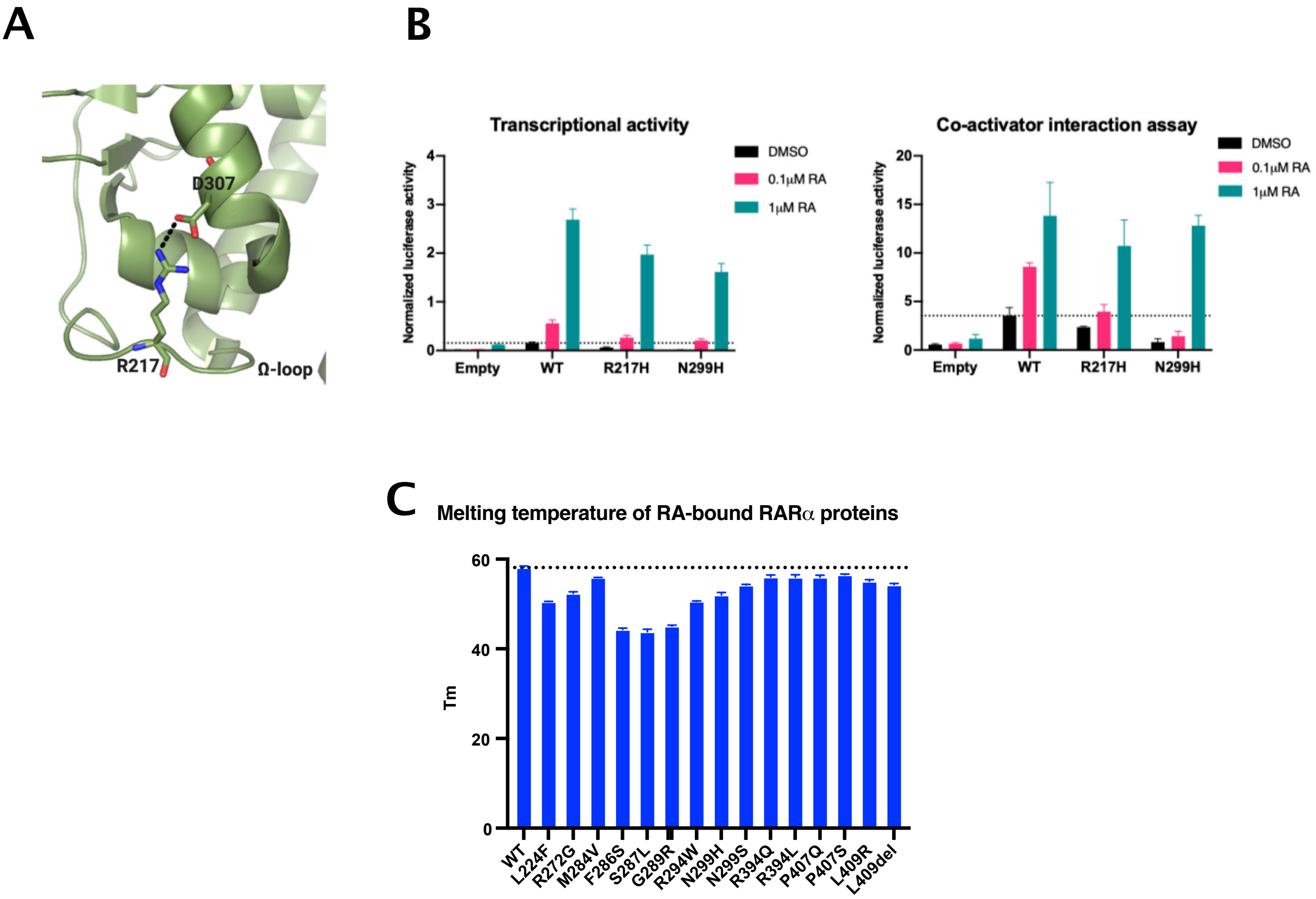


Supplementary Figure 5. RARA mutation R217H. (A) Location of R217 on omega loop. R217 resides on the omega-loop and forms polar contacts with D307. Mutation of this amino acid has been reported in APL patients who have developed RA treatment resistance. (B) Transcriptional activity and co-activator interactions of mutant RARα R217H and N299H. N=3, error bar = SD. (C) Melting temperatures of RA-bound WT and mutant RARα LBD proteins.
